# MMOD-induced structural changes of hydroxylase in soluble methane monooxygenase

**DOI:** 10.1101/331512

**Authors:** Hanseong Kim, Sojin An, Yeo Reum Park, Hara Jang, Sang Ho Park, Seung Jae Lee, Uhn-Soo Cho

## Abstract

Soluble methane monooxygenase in methanotrophs converts methane to methanol under ambient conditions^1-3^. The maximum catalytic activity of hydroxylase (MMOH) is achieved via interplay of its regulatory protein (MMOB) and reductase^4-6^. An additional auxiliary protein, MMOD, is believed to function as an inhibitor of the catalytic activity of MMOH; however, the mechanism of its action remains unknown^7,8^. Herein, we report the crystal structure of MMOH–MMOD complex from *Methylosinus sporium* strain 5 (2.6 Å), which illustrates that two molecules of MMOD associate symmetrically with the canyon region of MMOH in a manner similar to MMOB, indicating that MMOD competes with MMOB for MMOH recognition. Further, MMOD binding disrupts the geometry of the di-iron centre and opens the substrate access channel. Notably, the electron density of 1,6-hexanediol at the substrate access channel mimics products of sMMO in hydrocarbon oxidation. The crystal structure of MMOH–MMOD unravels the inhibitory mechanism by which MMOD suppresses the MMOH catalytic activity, and reveals how hydrocarbon substrates/products access to the di-iron centre.

---

Methanotrophs have an ability to use methane as their sole energy and carbon source^9,10^. The critical enzymes participating in this conversion process are particulate methane monooxygenase and soluble methane monooxygenase (sMMO)^11-13^. In particular, the sMMO enzyme, which belongs to the bacterial multicomponent monooxygenase superfamily, carries out the following reaction:

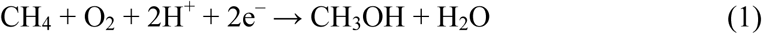

The effectiveness of catalysis by sMMO is known to be linked to the interplay of three sMMO components: MMO hydroxylase (MMOH), MMO reductase (MMOR) and MMO regulatory protein (MMOB)^4-6^. MMOH, which performs the chemical conversion of methane to methanol, is a homodimer of three subunits (α, β and γ), with a glutamate- and histidine-coordinated non-heme di-iron centre at each α-subunit^1,2^. MMOR shuttles electrons from nicotinamide adenine dinucleotide (NADH) to the di-iron centre in MMOH^14^. MMOB is a regulatory component that alters reduction potentials at the di-iron centre and helps the substrate to access the active site^15,16^. It has been shown that there is another auxiliary protein in the sMMO operon, called MMOD (*orfY*), which is proposed to inhibit the catalytic activity of MMOH^7,8^ and the ‘copper-switch’ in methanotrophs^17,18^. However, the molecular mechanism by which MMOD performs its function towards MMOH remains undetermined.

The crystal structure of MMOH and the NMR structures of MMOB and two MMOR domains have been determined, and their potential roles in the catalytic reaction have been proposed^19-23^. However, the absence of structures of MMOH in complex with its auxiliary proteins has made it difficult to fully elucidate the chemistry of sMMO in substrate oxidation. We reported previously the crystal structure of the MMOH–MMOB complex at 2.9 Å resolution^15^. Structural studies of MMOH–MMOB have addressed key questions about how MMOB modulates the geometry of the di-iron centre and the substrate access channel. However, to fully understand the enzymatic mechanism utilised for the conversion of methane to methanol by MMOH, additional structural information is required. In particular, structural information regarding how auxiliary proteins such as MMOR and MMOD influence the catalytic activity of MMOH upon binding, as well as the ingress and egress pathways of substrates and products in sMMO is currently missing.

We began to elucidate the molecular mechanism by which MMOD influences the catalytic activity of MMOH by determining the crystal structure of the MMOH–MMOD complex from *M. sporium* strain 5 at 2.6 Å resolution (Fig. 1a). The initial electron density map of the MMOH–MMOD complex was calculated by molecular replacement using *Methylosinus trichosporium* OB3b MMOH as the search model (PDB ID: 1MHY)^20^. *M. trichosporium* OB3b MMOH and *M. sporium* strain 5 MMOH enzymes are highly homologous, sharing primary sequence identity of 97% (α-subunit), 88% (β-subunit) and 87% (γ-subunit) (Extended Data Fig. 1a,b). The crystal structure of the MMOH–MMOD complex reveals that two copies of MMOD interact symmetrically with MMOH via its canyon region (Fig. 1a,b). The conserved core region of MMOD (residues 8–71) has a well-ordered structure and harbours a ββββα-fold architecture with four N-terminal anti-parallel β-strands and a long C-terminal α-helix (Fig. 1c and Extended Data Fig. 1c). Residues 1–11 and 76–111 of MMOD, which are less conserved among other strains, are disordered in the crystal structure.

**Figure 1.**
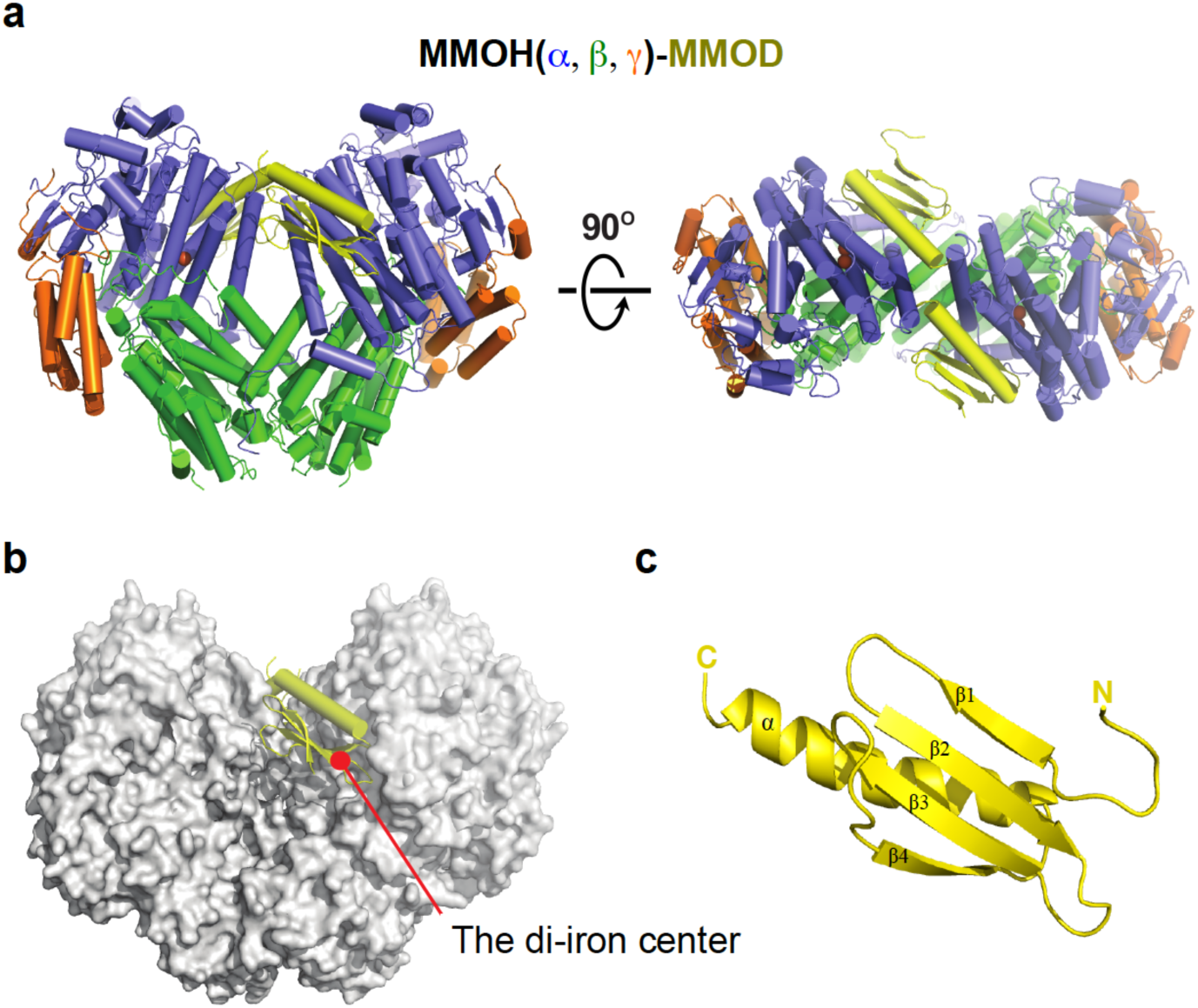
Crystal structure of the MMOH–MMOD complex. **a**, Front and top views of the MMOH–MMOD complex. MMOH is shown using a cartoon model (α-subunit: blue, β-subunit: green γ-subunit: orange); MMOD is coloured yellow. **b**, Front view of the MMOH–MMOD complex. MMOH is shown using a surface representation model (white) and MMOD (yellow) is shown in cartoon form. Red circle marks the location of the di-iron centre in MMOH. **c**, MMOD contains a ββββα-fold, which is shown in cartoon form. Illustrations of the protein structure used in all figures were generated with PyMOL (Delano Scientific, LLC).

MMOD binds to helices E, F and H of the MMOH α-subunit (MMOHα) through a β4 and an α1 helix (Fig. 2a,b). The side chain of Gln45 and the peptide backbone of residues 48 and 49 in MMOD β4 form hydrogen bond interactions with Thr241, Glu240 (helix F) and Asn214 (helix E) of MMOHα (Fig. 2b). Side chains and the backbone of MMOD Leu50 and Ser51 (MMOD loop) participate in hydrophobic and hydrogen bond interactions with Trp317 and Asp312 (helix H) of MMOHα (Fig. 2c). In the long α-helix of MMOD, residues including Ser54, His61, Ser64, His65, Arg68, Gln69 (hydrogen bond) and Val62 (hydrophobic) interact with Asp312 (helix H), Val218, Ser225 (helix E), Glu230 and Leu237 (helix F) of MMOHα (Fig. 2d).

**Figure 2.**
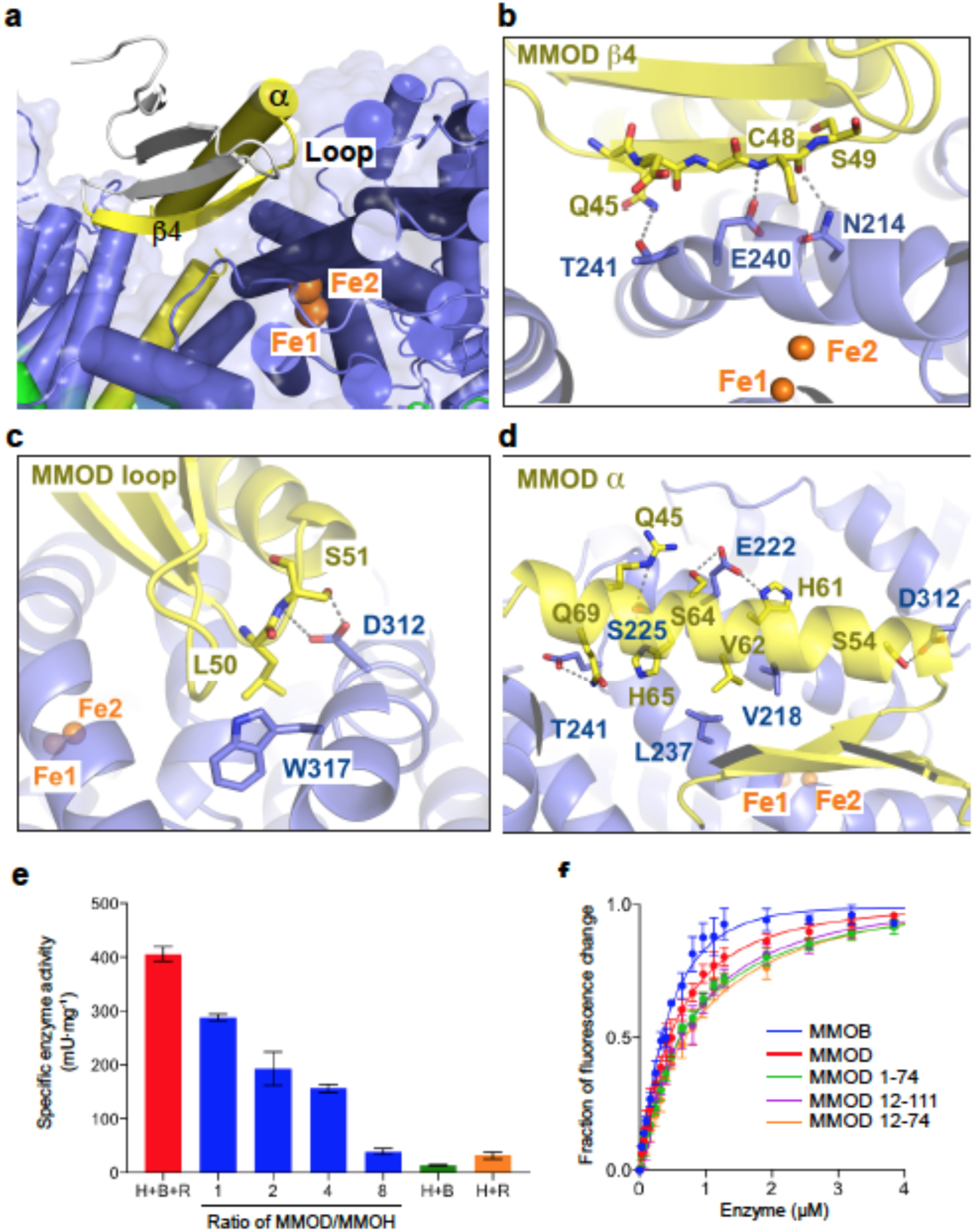
Key residues participate in both the MMOH–MMOD binding and the inhibitory activity of MMOD. **a**, MMOD β4, loop and helix α (yellow) directly participate in recognising the canyon region of MMOH. MMOD residues that do not interact with MMOH are coloured grey. The two iron atoms are labelled and coloured orange. **b–d**, Detailed interactions in the MMOH–MMOD complex. **e**, Enzymatic activities (n = 3) of MMOH in the presence of MMOB, MMOR and/or MMOD, as determined based on the conversion from propylene to propylene oxide in the presence of NADH. Condition H+B+R (red) indicates the absence of MMOD; blue indicates the addition of MMOD. Conditions H+B (green) and H+R (orange) represent the absence of MMOR and MMOB, respectively (n = 3, Ave ± SEM). **f**, Binding of MMOB, MMOD and truncated MMODs to MMOH as detected by fluorescence spectroscopy (n = 3). All error bars represent SEMs. Quenching of intrinsic fluorescence of MMOH was monitored by titration of the MMOB (blue), MMOD (red), 1–74 MMOD (green), 12–111 MMOD (purple), and 12–111 MMOD (orange). The *K*_D1_ and *K*_D2_ values were determined through nonlinear curve fitting for two binding sites of MMOH (0.32 μM).

The crystal structure of the MMOH–MMOB complex demonstrated that MMOB also associates with the canyon region of MMOH generated by two MMOH α-subunits (Extended Data Fig. 2a)^15^. Structural comparison between the MMOH–MMOB and MMOH–MMOD complexes indicates strongly that MMOD and MMOB compete for binding at the canyon region of MMOH, thereby inhibiting the catalytic activity of MMOH (Extended Data Fig. 2b). Indeed, the catalytic activity of MMOH is reduced in the presence of increasing amounts of MMOD, as reported earlier in *M. capsulatus* (Bath) sMMO counterparts (Fig. 2e)^7,8^. The affinity measurements of MMOB and MMOD towards MMOH using fluorescence quenching of MMOH demonstrated comparable dissociation constants (Fig. 2f and Extended Data Table 2). The truncated MMODs (residues 1–74, 12–111 and 12–74) displayed slightly, but not significantly, reduced binding affinities, suggesting that the conserved core region of MMOD observed in the crystal structure is essential for MMOH recognition (Extended Data Table 2).

In order to illustrate the influence of MMOD binding on the overall architecture of MMOH, as well as the geometry of the di-iron centre, the structures of MMOH and the MMOH–MMOD complex were compared. This comparison revealed a major conformational change in the N-terminus of the MMOH β-subunit (MMOHβ-NT) (Fig. 3a). Specifically, MMOHβ-NT becomes disordered upon MMOD binding (Extended Data Fig. 3a). A clash between the N-terminal helix of MMOHβ and the C-terminal long helix of MMOD results in MMOHβ-NT being pushed away, thereby dissociating it from MMOHα (Fig. 3b and Extended Data Fig. 3a). Since MMOHβ-NT functions as a ‘latch’ to hold the MMOHα (Extended Data Fig. 3a), the disorder of MMOHβ-NT (residues 1–56 of the β-subunit) initiates conformational changes in MMOHα, particularly helices A, B and C (Extended Data Fig. 3b, c). The di-iron centre at MMOHα is coordinated by four glutamates and two histidines provided by the four-helix bundle composed of helices B, C, E and F (Fig. 3c). The conformational changes in helices B and C, induced by MMOHβ-NT disorganisation, induce in turn geometric changes in the di-iron centre and rearrangement of the di-iron coordination (Fig. 3c,d). Particularly, His147 (helix C) displays a significant conformational change that dissociates it from Fe1 coordination and allows one water molecule to place in between Fe1 and His147 (Fig. 3d). The glutamate residues that coordinate the two iron atoms also rearrange their coordination upon MMOD association. For example, upon MMOD binding, both Glu114 and Glu144 recognise Fe1 in a bidentate/chelating manner through the carboxylate shift owing to the movement of helix B and C, respectively (Fig. 3c, d and Extended Data Fig. 3b, c). In addition, Glu243 interacts with Fe2 in a bidentate manner. To the best of our knowledge, the dissociation of His147 and bidentate coordination of Glu114 unravelled in the MMOH–MMOD complex have not been observed in previous MMOH-related structures (Fig. 3d and Extended Data Fig. 4). Notably, the discoordination of His147 from the di-iron active site (Fe1–His147 Nδ1 distance within two protomers: 5.27 Å / 4.98 Å) may further inhibit the catalytic activity of MMOH.

**Figure 3.**
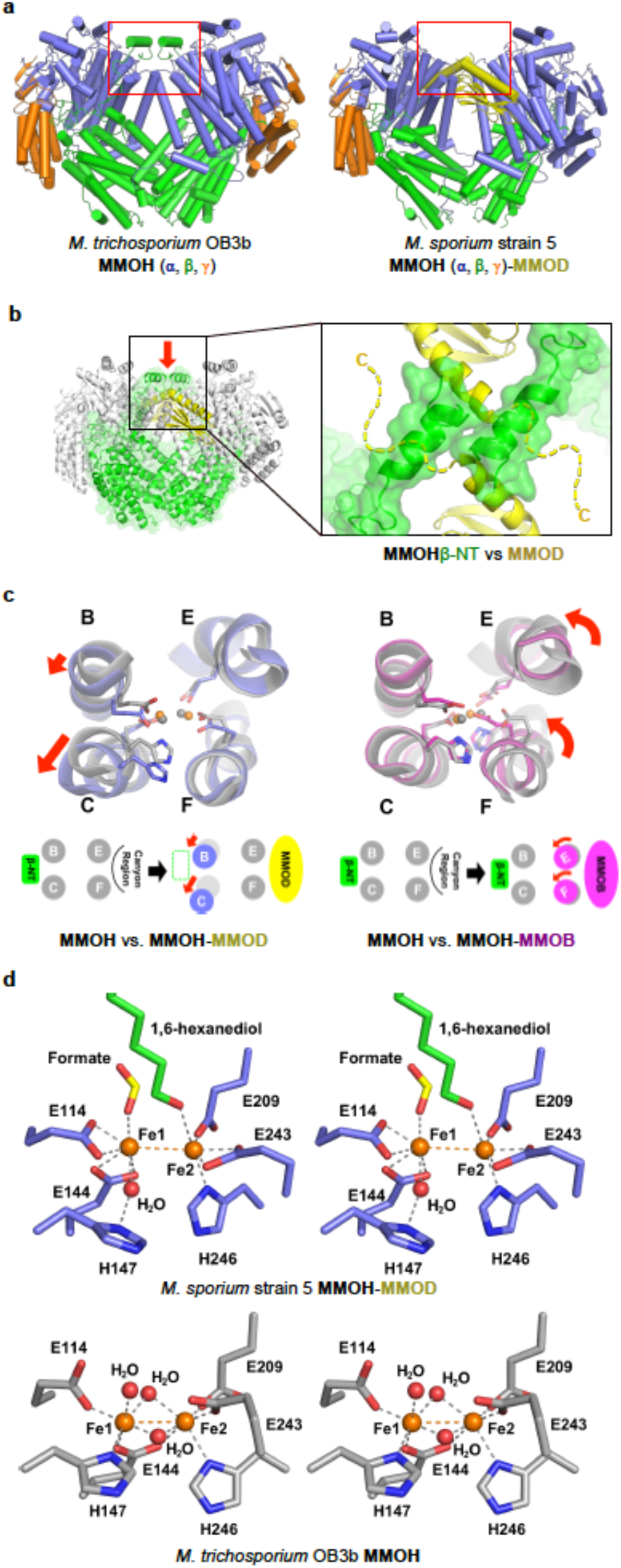
Structural comparison of MMOH (PDB ID: 1MHY)^20^ and MMOH–MMOD. **a**, Front views of MMOH and MMOH–MMOD. Red box indicates the region with the major structural differences. **b**, *M. sporium* strain 5 MMOD structures are overlaid onto *M. trichosporium* OB3b MMOH. N-terminus of MMOH β-subunit is displayed in green. MMOD C-terminal long helix is shown in yellow. Dashed yellow lines indicate the extended, disordered C-terminal region of MMOD. The red arrow indicates the viewing angle of inset. The inset shows the clash between MMOHβ-NT and the C-terminal helix of MMOD. **c**, Conformational changes in the MMOH four-helix bundle (helices B, C, E and F) upon MMOB and MMOD binding. Six residues that coordinate two iron atoms are on helices B, C, E and F. Red arrows indicate the translation and rotation of indicated helices upon MMOD and MMOB binding. **d**, Stereo-views of the di-iron centre geometry in MMOH–MMOD and MMOH. Two Fe atoms (orange) are surrounded and coordinated by four glutamate and two histidine residues. Water molecules (H_2_O) are displayed as red spheres.

The extent of structural reorganisation in the di-iron centre caused by MMOD binding is comparable to that caused by MMOB binding. It has been shown that the binding of MMOB induces conformational changes at helices E and F, which subsequently push Glu243 (helix F) into the di-iron centre (Fig. 3c and Extended Data Fig. 5a, b). Consequently, this change squeezes the two Fe atoms of the centre between Glu243 (helix F) and Glu144 (helix C), thereby leading to a change in reduction potential at the di-iron centre (Extended Data Fig. 5b)^16,24^. While MMOB binding induces the rotation of helices E and F, the MMOD binding disorganises MMOHβ-NT, thereby allowing the translational movement of helices B and C (Extended Data Figs. 3c and 5c). As a consequence, MMOB binding changes the reduction potential and MMOD binding inhibits the catalytic activity of MMOH.

Upon building the model structure of the MMOH–MMOD complex, we detected unidentified electron density near the di-iron centre. When we placed 1,6-hexanediol (one of the compounds used in the crystallisation) over this unknown electron density, a good fit was obtained after refinement (Fig. 4a). sMMO is known to oxidise a variety of hydrocarbons, including alkanes from methane to octane^5^. The presence of 1,6-hexanediol near the di-iron centre can be explained by opening of the channel, mediated by the side-chain rearrangement of Leu110 and Phe188, both of which function together as a gate for substrate and product passage to the active site (Fig. 4b)^15,19,25,26^. A similar type of channel opening has also been observed in the MMOH–MMOB complex (Extended Data Fig. 6a, b)^15^. This observation indicates that the channel opening in the MMOH–MMOD complex is also driven by the MMOD association. Examination of other previously reported product-soaked crystal structures of MMOH showed that products or product analogues are mostly trapped at cavities as a result of the channel closure mediated by Leu110 and Phe188 (Extended Data Fig. 6c). By contrast, 1,6-hexanediol is able to reach the di-iron centre via the substrate access channel in the MMOH–MMOD complex because the binding of MMOD opens the continuous channel (Fig. 4b). The disorganisation of MMOHβ-NT after the association of MMOH with MMOD, followed by structural relaxation of MMOHα generates additional tunnels inside of MMOH (Extended Data Fig. 7). The current MMOH–MMOD structure also supports the hypothesis that both substrate ingress and product egress take place through the substrate access channel, and not through the pore located near the active site, at least for hydrocarbon chain substrates such as hexane^5,26^.

**Figure 4.**
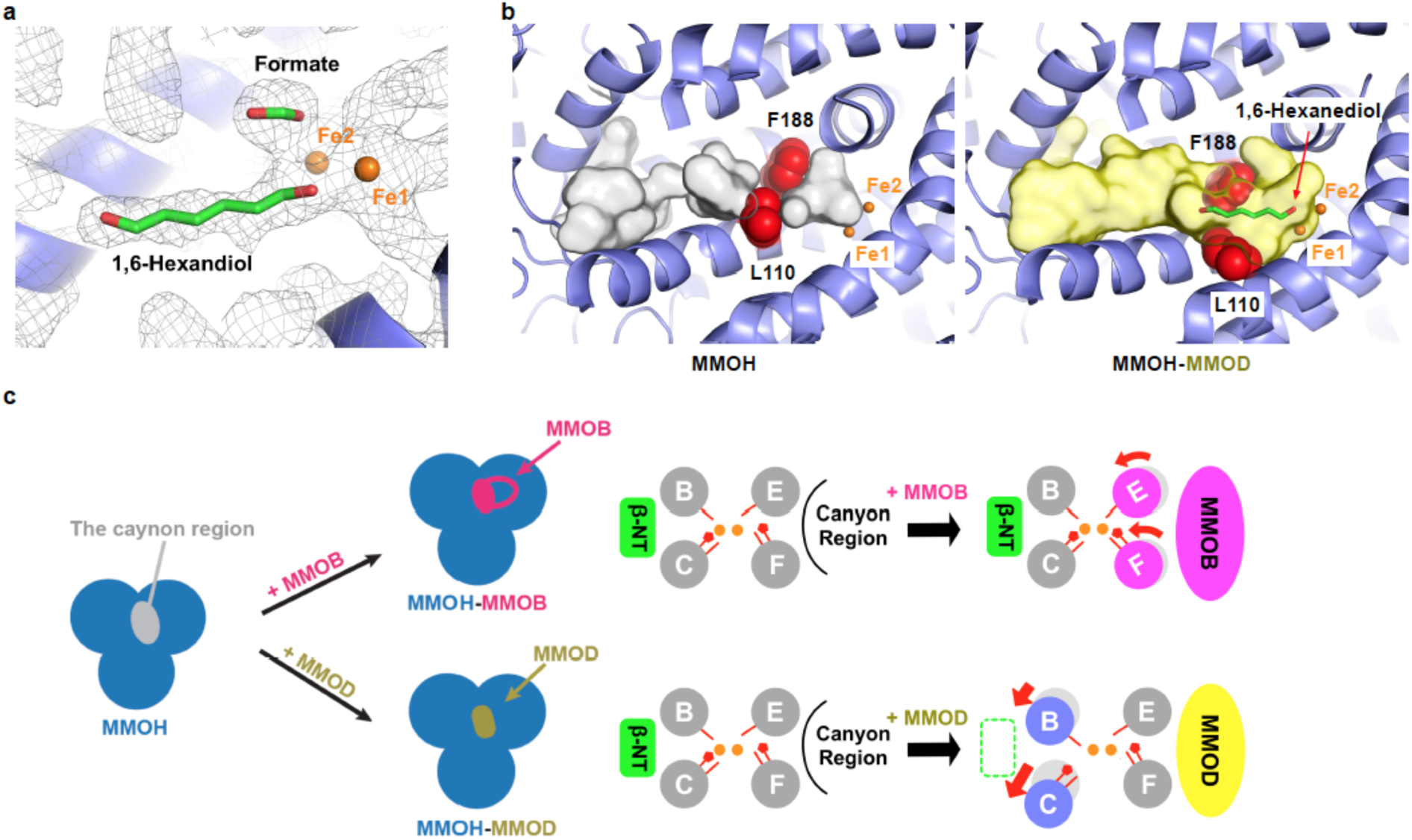
A product analogue (1,6-hexanediol) at the substrate access channel near the diiron centre and the channel opening in the MMOH–MMOD complex. **a**, The composite omit map (1.5-σ contour, calculated using the PHENIX software package^37^) of MMOH–MMOD near the di-iron centre. The electron density map is shown as mesh (grey), and 1,6-hexanediol and formate are coloured green. **b**, The substrate access channel in the *M. trichosporium* OB3b MMOH (grey, PDB ID: 1MHY)^20^ and the *M. sporium* strain 5 MMOH-MMOD complex (yellow). Two gate forming residues (Leu110 and Phe188) are shown as red spheres and 1,6-hexandiol is shown in green. The channels calculated using PyMOL are shown as surface models. **c**, A schematic model showing the association of MMOH with its auxiliary proteins– MMOB and MMOD, which compete for the canyon region of MMOH.

The crystal structures of the MMOH–MMOB and MMOH–MMOD complexes indicate that MMOB and MMOD associate competitively with the canyon region of MMOH. The results of hydrogen-deuterium exchange mass spectrometry analysis and computational docking studies in the MMOH–MMOR complex also corroborate the model in which MMOR associates with the canyon region of MMOH^27^. This strongly suggests that all three auxiliary proteins of sMMO modulate the catalytic activity of MMOH through their competitive association at the canyon region (Fig. 4c).

The MMOH–MMOD structure determined in this work indicates that MMOD functions as an inhibitor for MMOH by competing with MMOB for MMOH association, as well as by disrupting the productive geometry of the di-iron centre (Fig. 4c). The potential of MMOD to function as a ‘copper-switch’, in addition to its inhibitory role, needs to be investigated further^17,18^. Taken together, the crystal structure of the MMOH–MMOD complex and the MMOH–MMOB structure determined previously demonstrate the mechanism by which auxiliary proteins modulate the catalytic activity of MMO hydroxylase, which involves inducing differentiated conformational changes in MMOH as well as at the di-iron centre.

## Methods

### *M. sporium* strain 5 fermentation and purification of MMOH

*M. sporium* strain 5 (ATCC35069) was cultured in nitrate mineral salts (NMS) media (ATCC 1306) at 30 °C until OD_655_ reached 8–10 with a methane:air (v/v) ratio of 10–15%^28^. The cells were harvested by centrifugation (11,300 × *g*) for 20 min at 4°C. The cell pellets were suspended in a lysis buffer containing 3-(*N*-morpholino)propanesulfonic acid (MOPS, 25 mM), NaCl (25 mM), sodium thioglycolate (8 mM), _L_-cysteine (2 mM), (NH_4_)_2_Fe(SO4)_2_·6H_2_O (200 μM), MgCl_2_ (5 mM), DNase (0.25 μL/mL) and phenylmethane sulfonyl fluoride (PMSF, 0.04 mg/mL) at pH 6.5. The dissolved suspension was sonicated at 4 °C (CV334 model, Sonics), and the lysate was centrifuged at 30,000 × *g* for 45 min at 4 °C. The supernatant was carefully decanted and filtered through a 0.22-μm membrane (EMD Millipore). The filtrate was loaded onto DEAE Sepharose Fast Flow, Superdex 200, and Q Sepharose Fast Flow columns attached to the ATKA Pure 25L FPLC system (GE healthcare) to purify MMOH with >95% purity^4,29^. The purified MMOH was applied to a ferrozine assay to monitor the iron-ferrozine complex (562 nm), and the results revealed 3.9–4.2 Fe/MMOH with R^2^ values >0.999^15^.

### Expression and purification of MMOB and MMOR

Both recombinant *mmoB* (MMOB) and *mmoC* (MMOR) in a pET30a(+) plasmid were transformed into *Escherichia coli* BL21(DE3) cells (Novagen) and cultured in LB media containing 50 μg/mL kanamycin at 37 °C. The cultures were induced for 5 h with 1.0 mM isopropyl β-D-1-thiogalactopyranoside (IPTG) before harvesting by centrifugation at 4 °C. The MMOB cell pellets were re-suspended in solution containing phosphate (pH 6.0, 25 mM), NaCl (75 mM), MgCl_2_ (5 mM). Na_2_-EDTA (1 mM), DTT (dithiotheritol, 1 mM), DNase I (0.25 μL/mL) and PMSF (0.04 mg/mL). The suspended cells were lysed by sonication at 4 °C. The filtered supernatant was applied to Q Sepharose Fast Flow and Superdex 75 columns to purify MMOB with >95% purity^23,30^. The MMOR cell pellets were re-suspended in solution containing MOPS (pH 6.5, 25 mM), sodium thioglycolate (8 mM), _L_-cysteine (2 mM), NaCl (25 mM), (NH_4_)_2_Fe(SO4)_2_·6H_2_O (200 μM), MgCl_2_ (5 mM), DNase I (1 unit/mL) and PMSF (0.2 mM), and subsequently were sonicated for cell lysis. The lysed cells were centrifuged at 26,000 × *g* at 4 °C for 40 min, filtered (0.22-μm membrane filter) and loaded onto DEAE Sepharose Fast Flow and Q Sepharose columns to purify MMOR^31^. The ferrozine assay confirmed 2.1–2.2 Fe/MMOR with R^2^ values >0.999^29^.

### Activity measurement of MMOH

MMOH (1.0 μM), MMOB (2.0 μM) and MMOR (0.5 μM) were added to a 25-mM solution of MOPS (pH 7.5) containing DTT (1 mM) and NaCl (10 mM), and propylene gas was bubbled through the mixture (Hankook Special Gas) for 20 min. Subsequently, the mixture was incubated at 25 °C. Steady-state kinetics were measured using a Cary 60 UV-visible spectrometer by the addition of NADH at 340 nm (ε_340_ = 6,220 cm^−1^M^−1^). The inhibitory activities of MMOD were measured in the absence or presence of progressively increasing amounts of MMOD. Products were confirmed by gas chromatography (YL 6500GC system) using an Agilent HP-PLOT/Q stationery column (30 m × 0.535 mm × 40.00 μm). Independent experiments were repeated in triplicate to calculate the mean and the standard error of the mean (SEM).

### MMOD expression and purification

The N-terminal (His)_X6_ and maltose-binding protein (hisMBP)-tagged wild-type *mmoD* (MMOD) was prepared in *Escherichia coli* Rosetta (DE3) with auto-inducible media^32^. Cell pastes were re-suspended in solution containing Tris-HCl (pH 7.5, 30 mM), NaCl (500 mM) and β-mercaptoethanol (5 mM) with protease inhibitor cocktails. After sonication on ice for 3 min, soluble lysate was recovered by centrifugation at 20,000 × g for 30 min and was subsequently applied to a cobalt affinity resin (Takeda). The hisMBP-MMOD proteins were eluted with elution buffer (Tris-HCl = 30 mM, pH 7.5; NaCl = 500 mM; imidazole = 300 mM; β-mercaptoethanol = 5 mM). In order to remove the hisMBP-tag, the elution fraction and Tobacco Etch Virus protease (1:100 ratio) were mixed together in the amylose resin. After overnight incubation at 4 °C, the flow-through fraction, which contains MMOD, was collected from the amylose resin and loaded onto an HP Q column (GE Healthcare) equilibrated with HEPES (pH 7.5, 30 mM), NaCl (100 mM) and DTT (1 mM), and eluted in a 0 to 1000 mM NaCl gradient. Based on the results of SDS-PAGE gel electrophoresis, fractions containing MMOD were pooled. The pooled fractions were concentrated using Amicon (EMD Millipore) and applied to a Superdex200 (10/300) size exclusion chromatography (GE healthcare) column equilibrated in HEPES (pH 7.5, 30 mM), NaCl (100 mM) and TCEP (tris [2-carboxyethyl] phosphine, 1 mM).

### Crystallisation, data collection and structure determination

Purified MMOH and MMOD (in solution containing HEPES, 30 mM, pH 7.5; NaCl, 100 mM; and TCEP, 1 mM) were mixed in a 1:2 molar ratio and the final concentration was adjusted to 10 mg/mL. Plate shape crystals were grown within a week at room temperature using the hanging drop vapour diffusion method with a mixture of 10% (w/v) PEG 8000, 20% (v/v) ethylene glycol, 0.02 M 1,6-hexanediol, 0.02 M 1-butanol, 0.02 M 1,2-propanediol, 0.02 M 2-propanol, 0.02 M 1,4-butanediol, 0.02 M 1,3-propanediol, 0.89 M 1,3-butanediol, and 0.1 M MES/imidazole buffer (pH 6.5). After transferring to a cryo-protectant solution containing the precipitant and 35% (w/v) PEG 8000, crystals were flash-frozen in liquid nitrogen. The space group of the crystal was C2 with a = 179.8 Å, b = 125.8 Å, c = 126.4 Å, α = γ = 90° and β = 102.9°. The dataset was collected at Advanced Photon Source (APS) on beam line LS-CAT 21 ID-G. The 2.6 Å data were processed by HKL2000 (www.hkl-xray.com) and initial phases were calculated by molecular replacement (Phaser)^33^ using *Methylosinus trichosporium* OB3b MMOH as the search model (PDB ID: 1MHY)^20^. After molecular replacement, the electron density for MMOD was clearly visible. The MMOD model was subsequently manually built using COOT^34^ and the refinement was performed using PHENIX.refine^35^. Iron atoms and ligand restraints were generated using the program PHENIX (PHENIX.metal_coordination and PHENIX.ready_set) and applied during the refinement. No NCS restraints were applied during the refinement. The asymmetric unit contains two copies of each the MMOH α-subunit, β-subunit, γ-subunit and MMOD FL, and the R_work_ and R_free_ values of the final refined model were 17.8 and 22.6, respectively. Residues 1–11 (1–10) and 161–172 (159–171) of the MMOH α-subunit, residues 1–56 (1–56) of the MMOH β-subunit, residues 1–4 (1–2) of the MMOH γ-subunit and residues 1–11 (1–6) and 76–111 (75–111) of MMOD were disordered and invisible in the final model (Parenthesis indicates residues in the other MMOH protomer). Ramachandran analyses were performed by MolProbity^36^ and the results were as follows: 95.4 (favoured), 4.5 (allowed) and 0.1% (outlier).

### Binding affinity measurements

The fluorescence spectra were collected at 25 °C using the PTI QM-400 (HORIBA, CANADA) with monochromators for both excitation and emission^24^. Fluorescence spectra of MMOH (0.32 μM of MMOH in 300 μL of 25 mM MOPS and 1 mM DTT [pH 7.0]) were determined using an excitation wavelength of 282 nm. Fluorescence was quantified by integration of fluorescence emission bands with a maximum at 363 nm. The MMOH fluorescence was quenched by titration with MMOB, MMOD and truncated MMODs (residues 1–74, 12–111 and 12–74). These enzymes were titrated to MMOH in the concentration range from 0 to 15 μM. Dissociation constants were determined by fitting the intensity of emission at 363 nm (Origin2018b and Wolfram Mathmatica). Independent experiments were repeated in triplicate to allow the calculation of the average ± SEM.

## Acknowledgments

We thank the staff at the Advanced Photon Source LS-CAT beamline for their advice and assistance with data collection. We thank Dr Stephen Ragsdale, Dr Youngmin You and Dr David Ballou for helpful discussions and comments on the manuscript. This work was supported in part by grants (the C1 Gas Refinery Program through the National Research Foundation of Korea [NRF-2015M3D3A1A01064876]) to S.J.L., and by grants (The Biomedical Research Council [BMRC] bridging fund [University of Michigan] and R01 DK111465) to U.S.C.

## Author Contributions

H.K. obtained the crystals, solved, and refined the structures, analysed data and wrote the manuscript; S.A. purified proteins, analysed data and wrote the manuscript; Y.R.P. cultured *M. sporium* 5 and purified proteins for catalytic activity and affinity measurements; H.J. expressed and purified truncated MMOD; S.H.P. assisted structural determination; S.J.L. designed experiments for catalytic activity and binding measurement, analysed data and wrote the manuscript; and U.S.C. directed the project, designed experiments, analysed data and wrote the manuscript. All authors discussed the results and commented on the manuscript.

## Author Information

### Data availability

Coordinates and structure factors for the reported crystal structure of the *M. sporium* strain 5 MMOH–MMOD complex have been deposited with the Protein Data Bank under accession code 6D7K.

## Competing interests

The authors declare no competing interests.

**Extended Data Table 1.** | Data collection and refinement statistics.

| *M. sporium* strain 5 MMOH–MMOD Complex* |  |
| --- | --- |
| Data collection |  |
| Space group | C 1 2 1 |
| Cell dimensions |  |
| a, b, c (Å) | 179.78, 125.78, 126.38 |
| α, β, γ (°) | 90.00, 102.89, 90.00 |
| Resolution (Å) | 50.00–2.60 (2.64–2.60) ** |
| R<sub>sym</sub> or R<sub>merge</sub> | 12.4 (79.3) |
| I / σI | 12.8 (1.6) |
| Completeness (%) | 99.5 (98.9) |
| Redundancy | 4.3 (4.2) |
| Number of reflections | 362,653 |
| Unique | 84,164 |
| Refinement |  |
| Resolution (Å) | 22.29–2.60 (2.66–2.60) |
| No. reflections | 83,613 |
| R<sub>work</sub> / R<sub>free</sub> | 0.179 / 0.220 |
| No. of atoms |  |
| Protein | 17,401 |
| Ligand/ion | 28 |
| Water | 339 |
| B-factors |  |
| Protein | 48.71 |
| Ligand/ion | 38.88 |
| Water | 42.40 |
| R.M.S. deviations |  |
| Bond lengths (Å) | 0.008 |
| Bond angles (°) | 0.891 |

\*PDB ID 6D7K \*\*Values in parentheses are for highest-resolution shell

**Extended Data Table 2.** | Dissociation constants for binding of MMOH with MMOB, MMOD, and truncated MMODs.

| Enzymes | $K_{D1}$ ($\mu\text{M}$) | $K_{D2}$ ($\mu\text{M}$) |
| --- | --- | --- |
| MMOB | $0.04 \pm 0.01$ | $0.86 \pm 0.16$ |
| MMOD | $0.59 \pm 0.03$ | $0.90 \pm 0.14$ |
| MMOD (1–74) | $0.82 \pm 0.10$ | $2.69 \pm 0.28$ |
| MMOD (12–111) | $0.93 \pm 0.06$ | $2.75 \pm 0.26$ |
| MMOD (12–74) | $1.14 \pm 0.10$ | $2.90 \pm 0.61$ |

The intensities of intrinsic fluorescence of MMOH were partially quenched by titration of enzymes, and two binding sites of MMOH were used with nonlinear curve fitting (equation 1, $n = 3$ , Ave ± SEM). E represents MMOB, MMOD, and truncated MMOD (1–74, 12–111 and 12–74).

**Extended Data Figure 1.**
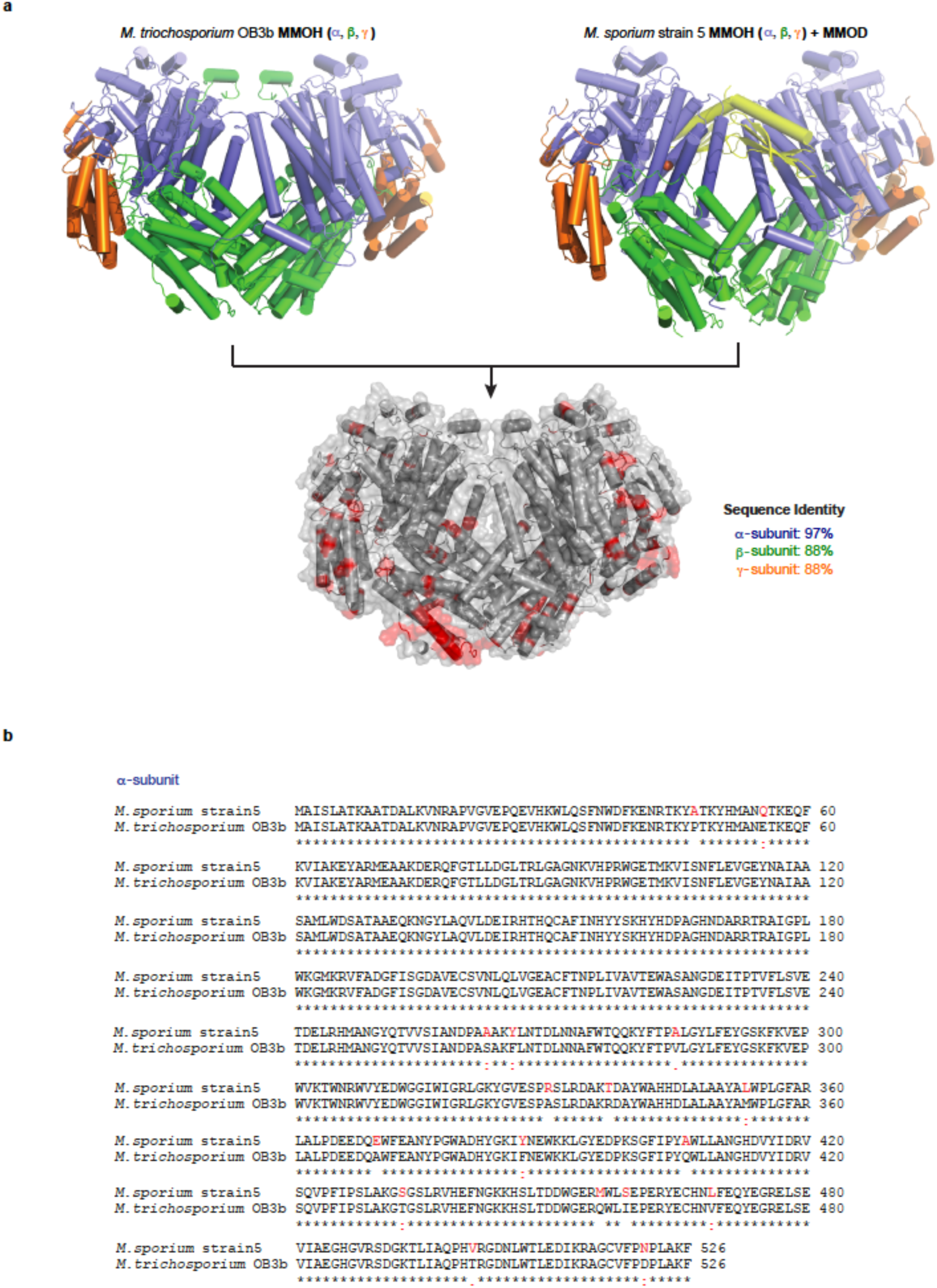
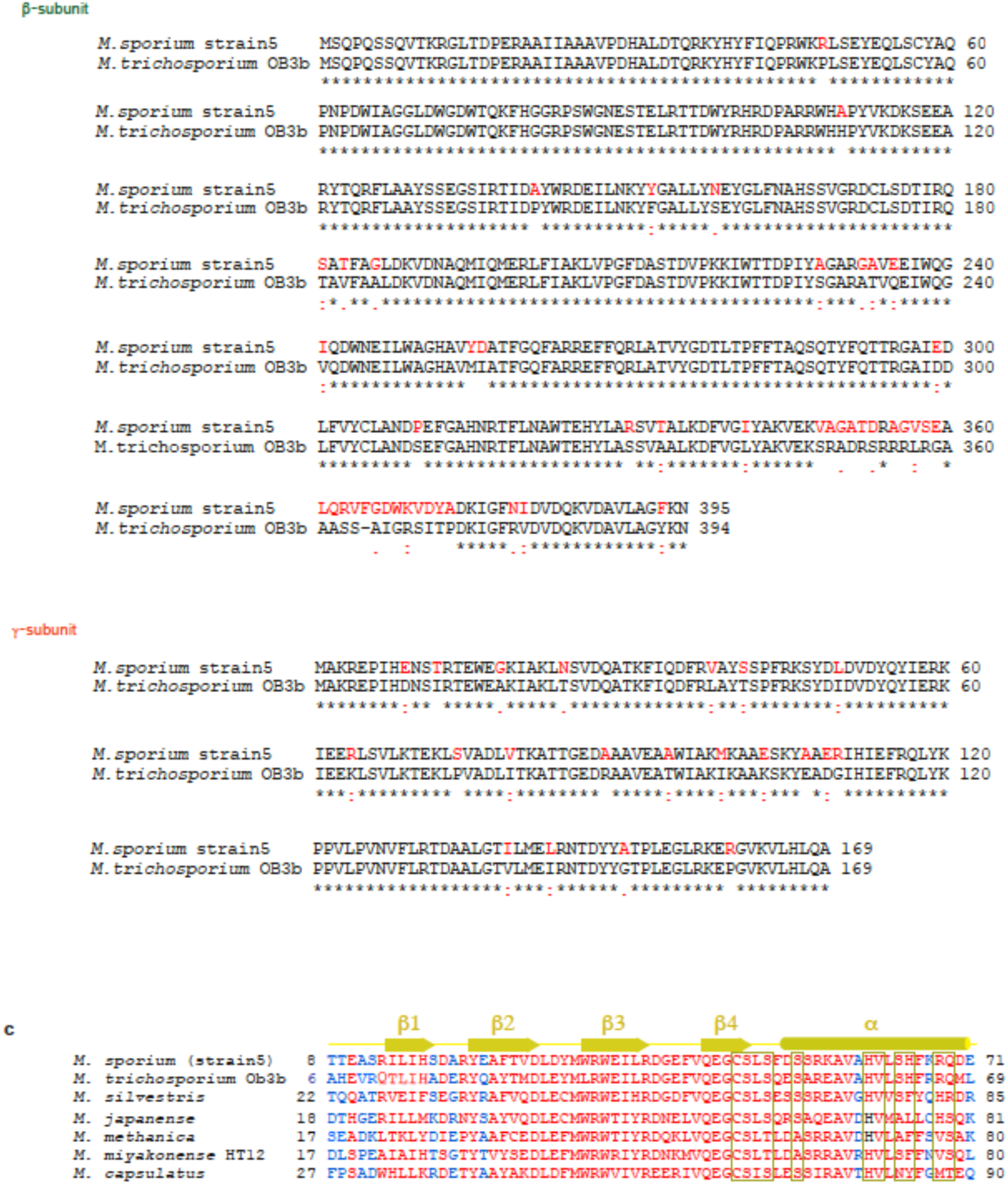
Primary sequence conservation of *M. trichosporium* OB3b and *M. sporium* strain 5 MMOHs and multiple sequence alignment of MMOD. **a**, Structure comparison of *M. trichosporium* OB3b MMOH and *M. sporium* strain 5 MMOH–MMOD. The primary sequence identities of α-, β- and γ-subunits in *M. trichosporium* OB3b and the *M. sporium* strain 5 MMOHs are 97%, 88% and 88%, respectively. Residues that differ in *M. trichosporium* OB3b and the *M. sporium* strain 5 MMOHs are coloured red. Residues involved in MMOD recognition and di-iron centre formation are 100% conserved in *M. trichosporium* OB3b and the *M. sporium* strain 5 MMOHs. **b**, The primary sequence comparison of α-, β- and γ-subunits in *M. trichosporium* OB3b and the *M. sporium* strain 5 MMOHs. MMOH residues that differ in *M. trichosporium* OB3b and the *M. sporium* strain 5 are coloured red. **c**, The multiple sequence alignment of the MMOD core region (residues 8–71 in *M. sporium* strain 5 MMOD) among methanotrophs. Residues that have high sequence similarity are coloured red; others are coloured blue. Residues that participate in the MMOH interactions are boxed in yellow. Corresponding secondary structures of MMOD are indicated above the sequence alignment.

**Extended Data Figure 2.**
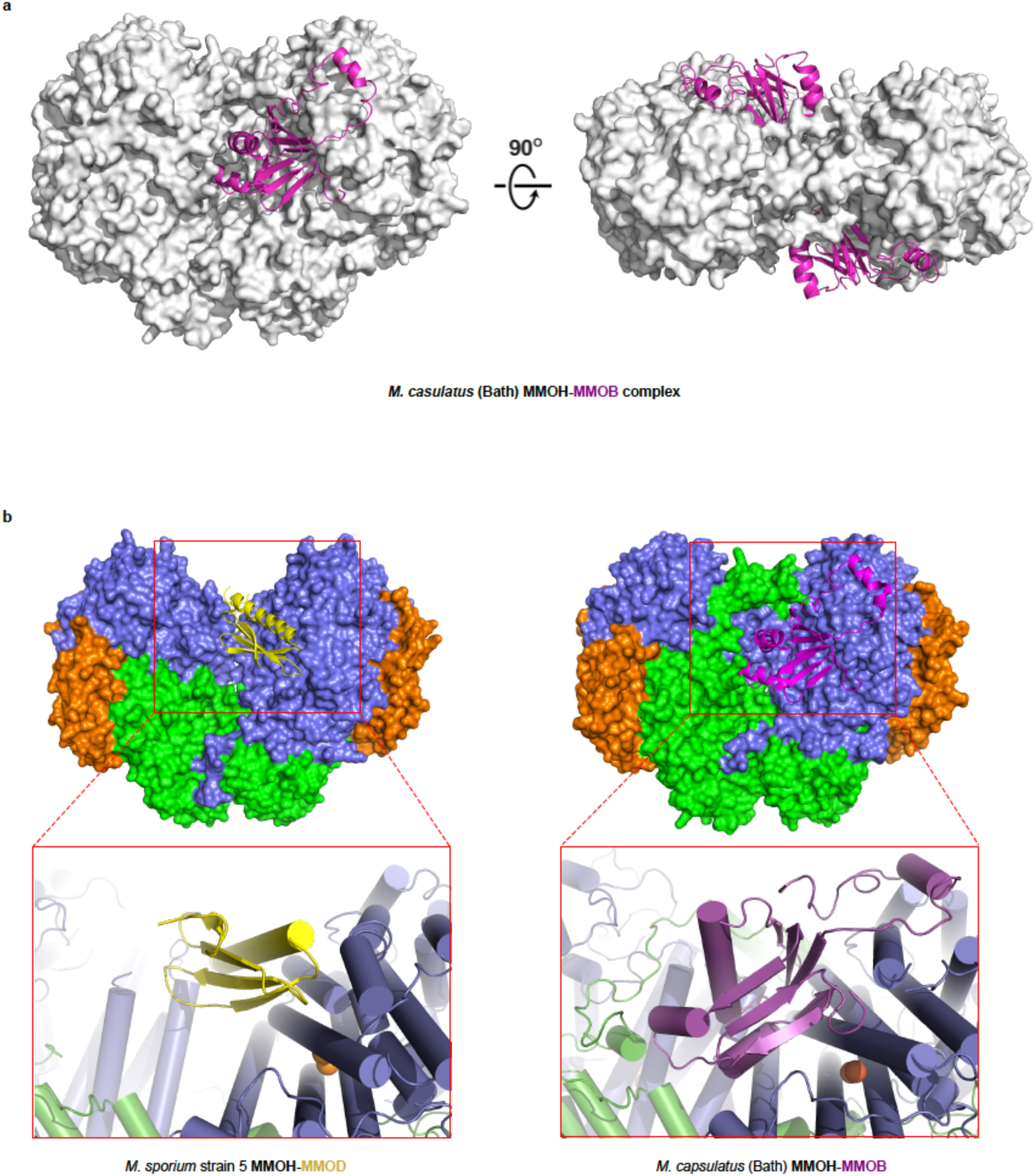
Crystal structure of the *M. capulatus* MMOH–MMOB complex (PDB ID: 4GAM)^15^ and the structural comparison of MMOD (*M. sporium* strain 5) and MMOB (*M. capsulatus*) binding to MMOH. **a**, front and top views of the *M. capsulatus* MMOH–MMOB complex. MMOH is shown using surface representation model (white) and MMOB (purple) is shown using a cartoon model. **b**, The structural comparison of MMOH– MMOD (*M. sporium* strain 5) and MMOH–MMOB (*M. capsulatus*) complexes. Areas with the red box are magnified to visualise the binding patterns of MMOD (yellow) and MMOB (purple) at the canyon region of MMOH. Although the overall architectures of MMOD and MMOB are quite distinct, both MMOD and MMOB bind competitively at the same canyon region of MMOH.

**Extended Data Figure 3.**
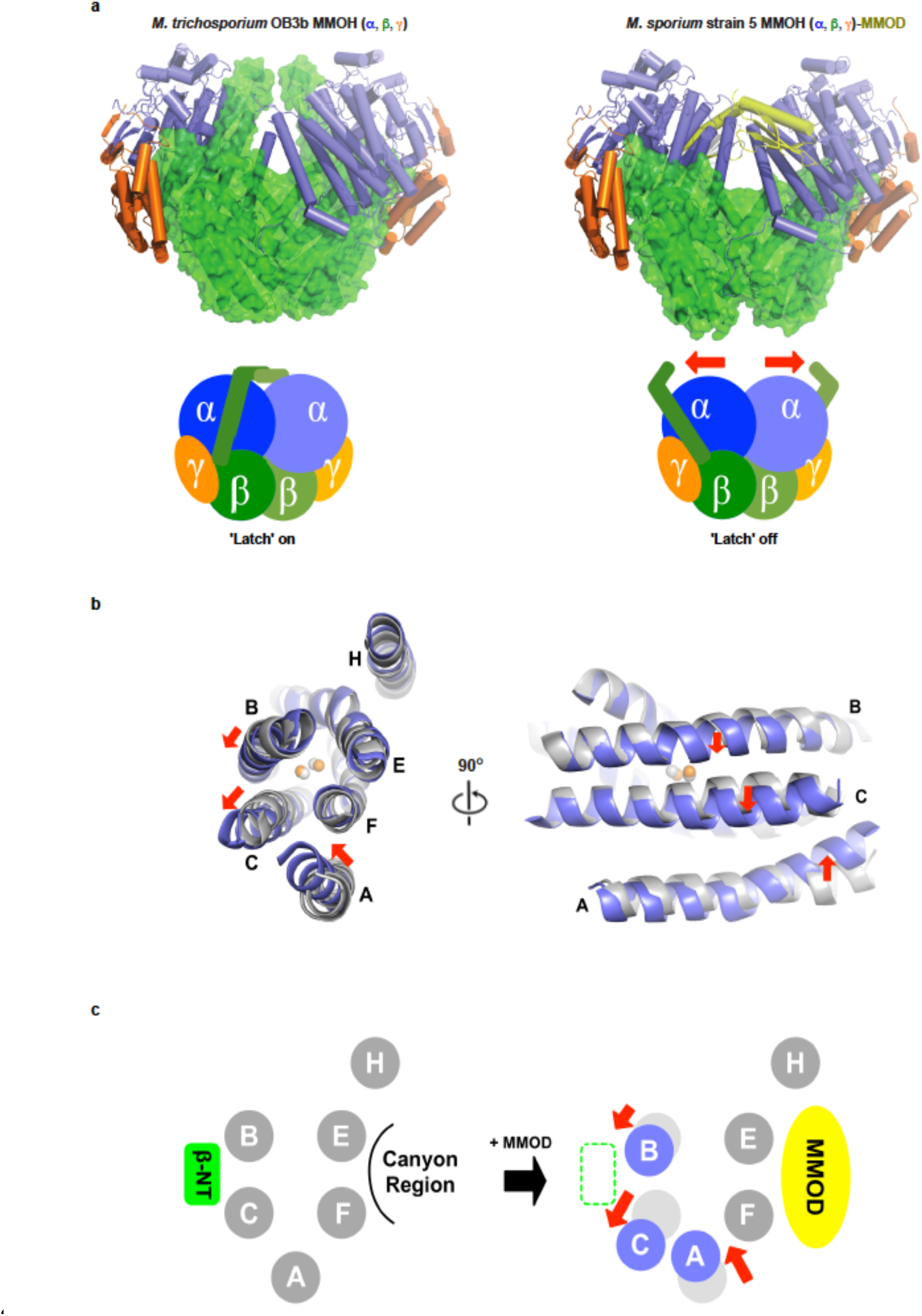
Conformational changes in MMOH upon MMOD binding. **a**, Structural comparison of MMOH (*M. trichosporium* OB3b) and the MMOH–MMOD complex (*M. sporium* strain 5). MMOHβ is shown using the surface representation model and other subunits (α and γ), including MMOD, are shown as cartoon models. Structural comparison indicates that the N-terminal region of MMOHβ (β-NT) is disordered upon MMOD association. The schematic model shows that MMOHβ-NT may function as a ‘latch’ to tightly lock MMOHα helix bundles. MMOD binding unlocks the ‘latch’ and relaxes the overall helix architecture of MMOHα, as shown in **b** and **c**. **b**, Top and side views of MMOHα helices (A, B, C, E, F and H) before (grey, PDB ID: 1MHY)^20^ and after (blue) MMOD association. MMOD binding breaks the interaction between MMOHβ-NT (‘latch off’) of two protomers and thus induces conformational relaxation of MMOHα, including helices A, B, and C. Red arrows indicate the direction of helix movement after MMOD association and the MMOHβ-NT latch off. Two iron atoms are shown using the sphere model (before: grey, after: orange)**. c**, Schematic representation of MMOHα helix movement upon MMOD association. MMOD binding causes MMOHβ-NT to ‘latch off’, which then leads to the movement of helices A, B and C owing to the loss of support from MMOHβ-NT.

**Extended Data Figure 4.**
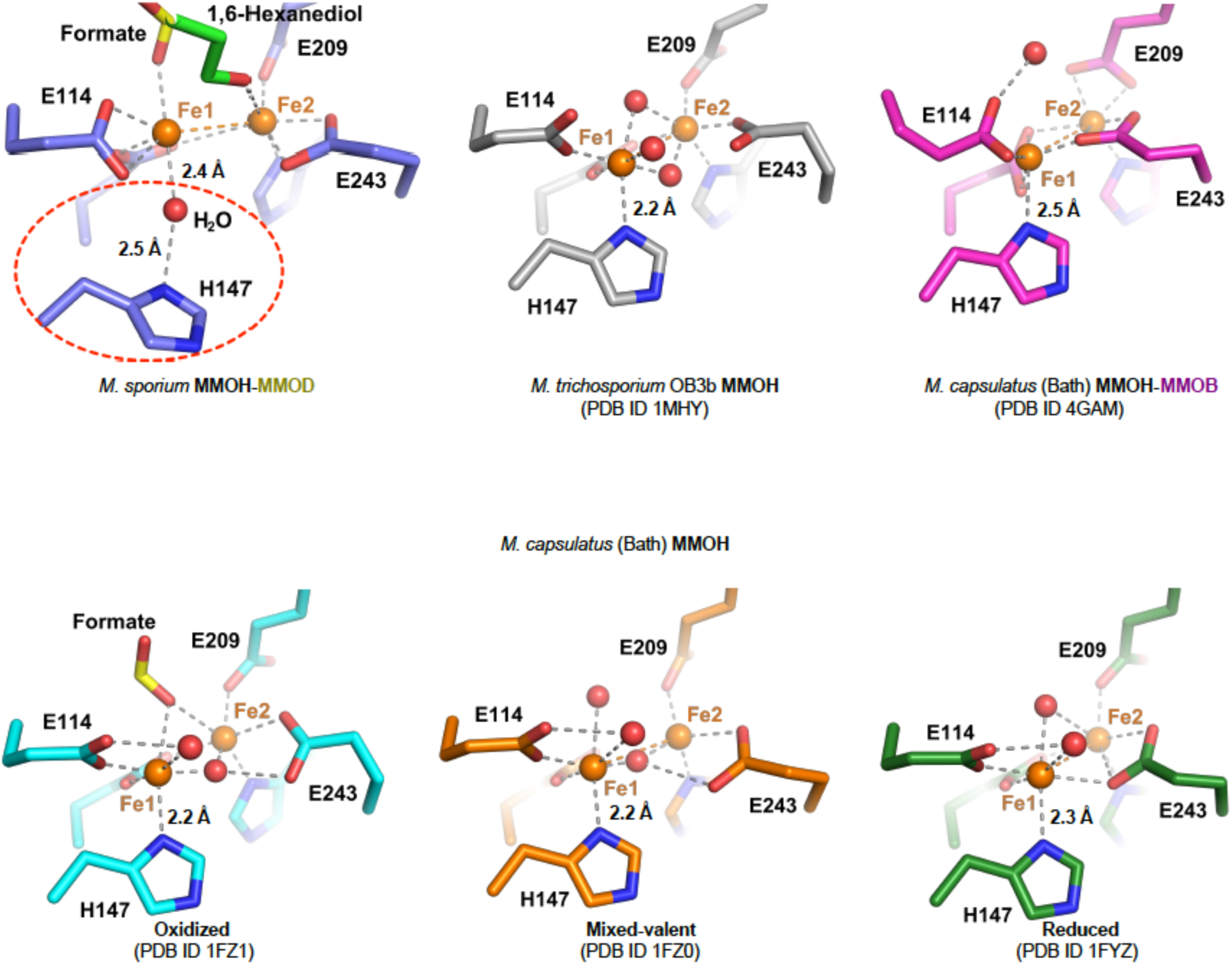
Geometry of the di-iron active site in the MMOH–MMOD complex and its structural comparison with other known MMOH structures. The geometry of the di-iron centre in the MMOH–MMOD complex differs significantly from other known MMOH di-iron centre geometries. There is dissociation of His147 from Fe1 coordination (red dotted circle) as well as bidentate recognition of Glu114 towards Fe1. A water molecule bridges the Fe1 coordination of His147. Distances between Fe1 and Nδ1 atom of His147 are indicated in each structure.

**Extended Data Figure 5.**
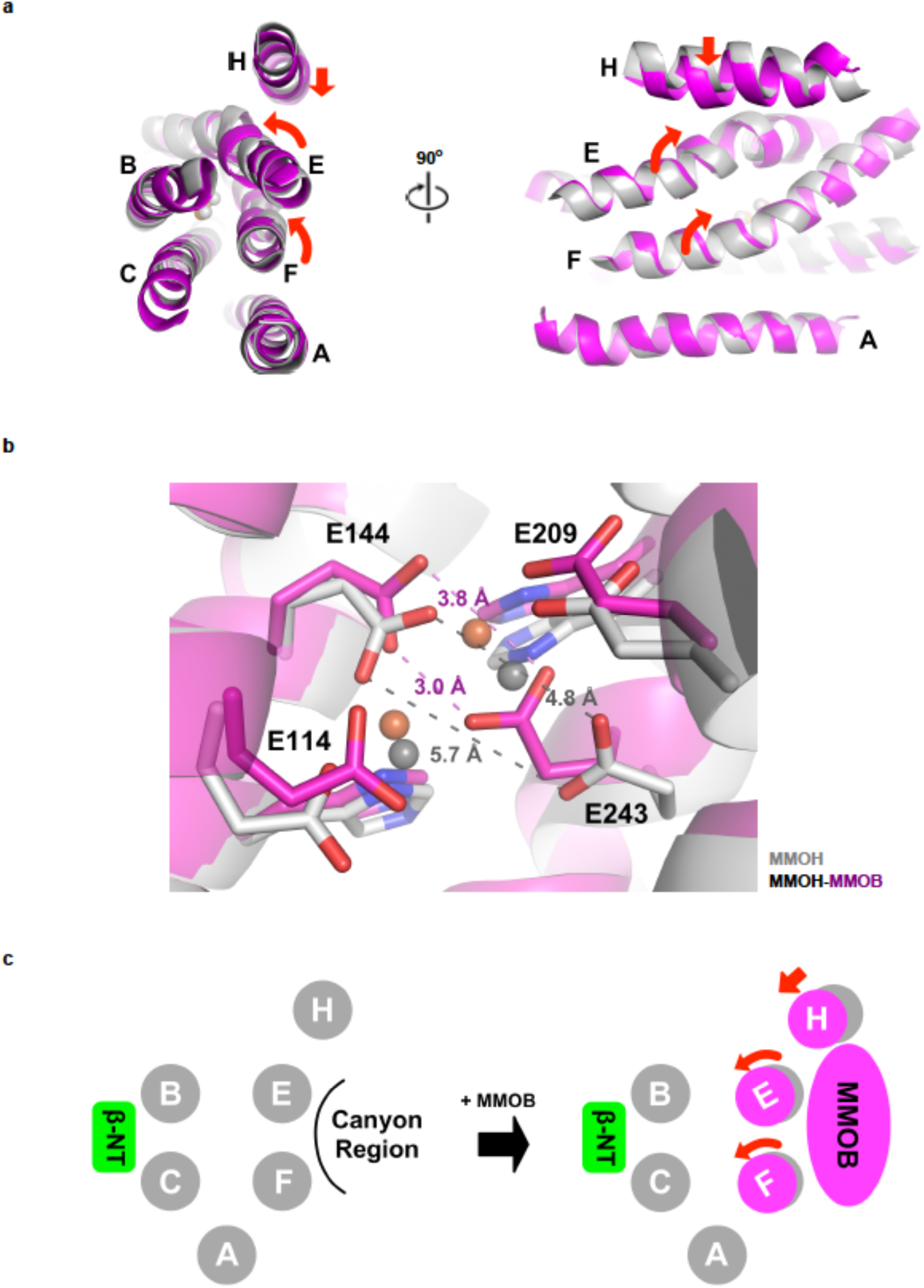
The movement of helices in MMOHα upon MMOB binding. **a**, Top and side-views of MMOHα helices (A, B, C, E, F and H) before (grey, PDB ID: 1FZ1)^38^ and after (purple, PDB ID: 4GAM)^15^ MMOB association in the *M. capsulatus* (Bath) MMOH– MMOB complex. MMOB binding causes the movement of helix H and the counter-clockwise rotation of helices E and F in MMOHα. **b**, Geometry changes in the di-iron centre upon MMOB association. MMOB binding induces the rotation of helices E and F, which contain iron coordinating Glu209 (helix E) as well as Glu243 and His246 (helix F). Particularly, rotation of helix F forces Glu243 to get close to the di-iron centre, thereby decreasing the distance between Glu144 and Glu243 (oxygen–oxygen distances decrease from 5.7 Å and 4.8 Å to 3.0 ± 0.2 Å and 3.8 ± 0.2 Å). Consequently, two irons become tightly coordinated by Glu144 and Glu243, causing changes in the reduction potential. **c**, The schematic representation of MMOHα helix movement upon MMOB association. MMOB binding causes the rotations of MMOHα helices E and F, which induce the rearrangement of the geometry of the di-iron centre.

**Extended Data Figure 6.**
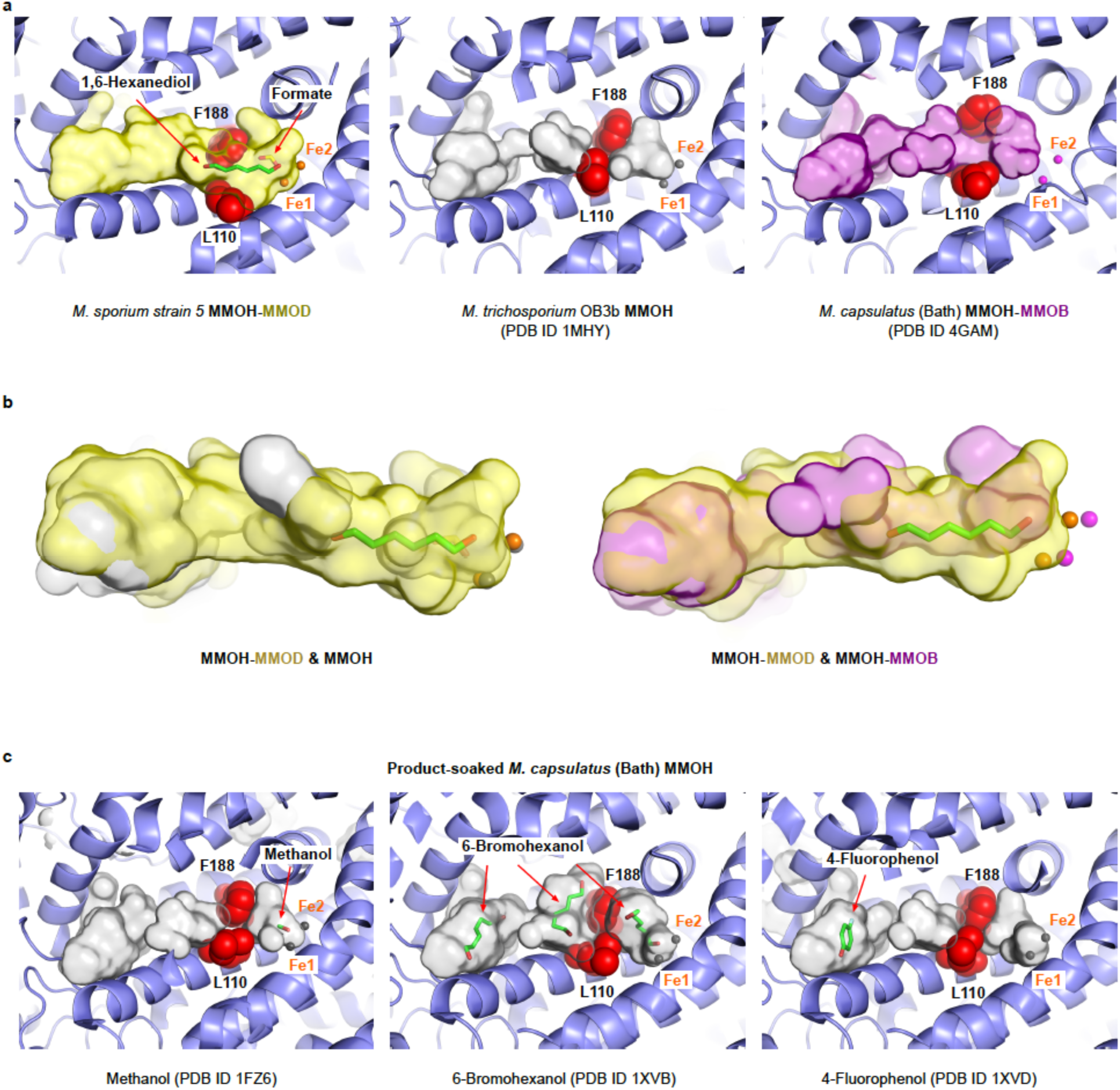
Comparison of the substrate access channel in MMOH (PDB ID: 1MHY)^20^, MMOH–MMOB (PDB ID: 4GAM)^15^ and MMOH–MMOD. **a**, Cavities of the substrate access channel in MMOH (grey), MMOH–MMOB (purple) and MMOH–MMOD (yellow) are displayed as surface representation models. The gate forming residues (Leu110 and Phe188) are shown as spheres of red colour. A product analogue, 1,6-hexanediol, is also shown using stick representation (green). In MMOH, the substrate access channel to the di-iron centre is disconnected by gate-forming residues. In both MMOH–MMOB and MMOH–MMOD, the gate is open. **b**, Comparison of the substrate access channel in MMOH–MMOD and MMOH, as well as MMOH–MMOD and MMOH–MMOB. In both cases, the substrate access channel in the MMOH-MMOD complex is wider, probably owing to the overall structural relaxation of MMOHα caused by MMOHβ-NT disorganisation. **c**, The substrate access channel in *M. capsulatus* (Bath) MMOH soaked with various products or product analogues. Substrate access channels in the presence of methanol (PDB ID: 1FZ6)^39^, 6-bromohexanol (PDB ID: 1XVB)^26^, and 4-florophenol (PDB ID: 1XVD)^26^ are displayed as surface representation models. In all cases, gate closure is mediated by Leu110 and Phe188, and products (substrate analogues) are trapped in either one or all of the three cavities. All channels are shown as surface models, which were calculated using PyMOL.

**Extended Data Figure 7.**
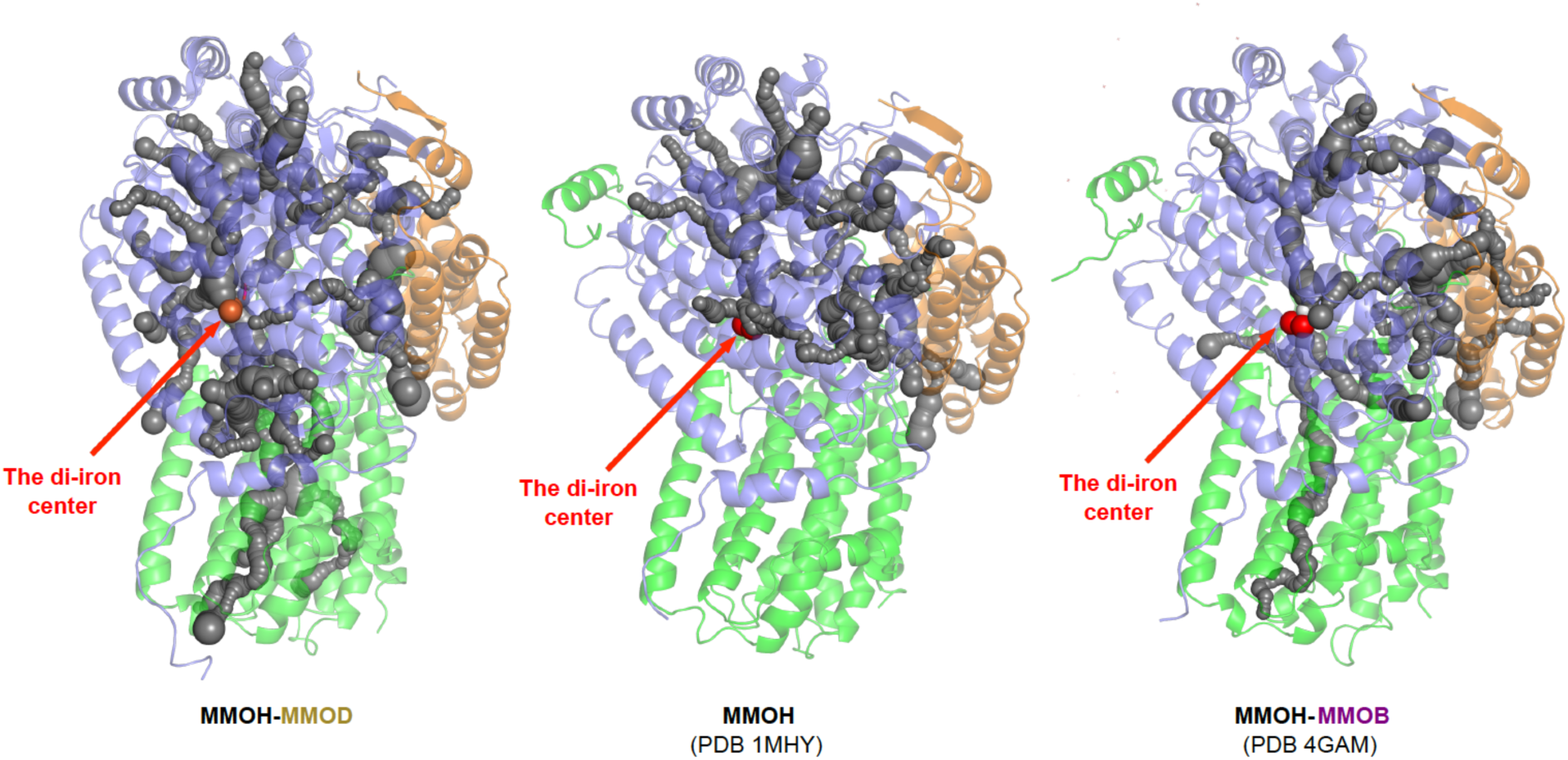
Comparisons of internal cavities within the MMOH protomer of MMOH–MMOD, MMOH and MMOH–MMOB. The MMOH protomer is presented in cartoon form; α-subunit (blue), β-subunit (green) and γ-subunit (orange). The di-iron centre is shown as orange (MMOH–MMOD) and red spheres (MMOH and MMOH–MMOB). Internal cavities calculated using CAVER^40^ are shown as chains of grey spheres. The CAVER analysis of MMOH protomers reveals that MMOD association produces more internal cavities when compared to MMOH alone and MMOH–MMOB. CAVER 3.0 (PyMOL plug-in) with the default setting was used to calculate internal cavities.

